# Exploring the role of abiotic factors on the seedling recruitment of three plant species in the Colombian Amazon

**DOI:** 10.1101/2022.07.30.502172

**Authors:** Ana C Palma, Pablo R. Stevenson

**Author notes:** Corresponding Author: Ana C. Palma, Address: Centro de Investigaciones Ecológicas Macarena. Universidad de Los Andes, Cra. 1 #18a-12, Bogotá, Cundinamarca, Colombia.

## Abstract

Our work seeks to explore certain abiotic factors that could affect seedling recruitment in neotropical moist forest. We selected three plant species (*Caraipa* sp., *Tachigali* sp., and *Dicranostyles* sp.) and investigated the role of nutrients, proximity to the parent tree, competition (intra-and inter-specific) and light availability on seedling growth and survival. We planted seedlings in the forest understory and in a shade house and applied four different treatments to evaluate the role of abiotic factors on recruitment. Our results show that the studied species were not negatively influenced by distance-dependent processes or by competition during the study period (6-7 mo.). We found that light availability was the most important factor determining seedlings’ early performance. The studied species showed distinct responses to the different treatments; therefore, it is hard to draw general conclusions about which factors might be affecting seedling recruitment and promoting the high plant diversity of tropical moist forest.

## Introduction

Processes involved in plant recruitment have been studied extensively to understand the critical role of early developmental stages in plant population dynamics (Howe & Smallwood 1982; Schupp & Fuentes 1995; Nathan & Muller-Landau 2000). These processes are of interest because of their implications explaining the rich diversity of plant communities (Janzen 1970; Hubbell 1980; Harms et al. 2000). Janzen (1970) and Connell (1971) suggest that the high density of seeds and seedlings under parent trees attracts predators, which decreases the probability of survival of seeds and seedlings. According to the Janzen-Connell theory, under parent trees and conspecifics there can be greater intra-specific competition and a greater probability of contagion of pathogens between seedlings (Connell 1971; Augspurger 1983). This high mortality near conspecifics could favour the high diversity of plant communities since the seeds of less abundant species could have a greater probability of regeneration.

Recruitment patterns are attributed to a wide range of factors, their interactions and variations in time and space (Nathan & Casagrandi 2004). Although seedling survival depends on both biotic and abiotic factors, biotic factors (e.g., herbivory, pathogens) have been studied further (Janzen 1970; Connell 1971; Augspurger 1983; Shupp 1990; Packer & Clay 2003; Bagchi et al. 2014; among others), overlooking the potential effects of abiotic factors. While some studies evaluated the effect of various abiotic factors such as light and water availability, litter accumulation or available area (Kenkel 1988; Mori & Mizumachi 2005; Messaoud & Houle 2006; Mithen et al. 1984), few studies have focused on soil characteristics and how these may affect seedling recruitment (but see Challinor 1968; Antonovics & Levin 1980; Mori & Mizumachi 2005; Maestre et al. 2005; Maestre & Cortina 2004). Most of these studies have been carried out in temperate forests, in disturbed areas (e.g., Heydari et al 2017) and only a few studies have addressed how different soil types, water availability, light or nutrients can affect the growth of seedlings in neotropical moist forests (i.e., Coomes & Grubb, 1998; Montgomery & Chazdon, 2002; Baraloto et al., 2005 and 2006; Fine et al., 2004, Anderson, 2009 and Umaña et al, 2018).

The general explanation for the poor success of seeds and seedlings near parent trees centres on the role of predators, pathogens, and competition due to higher densities (Janzen 1970; Connell 1971; Augspurger 1983; Schupp 1990). However, the role of the parent tree and its possible abiotic effects on the success of seedlings has not been widely evaluated. Abiotic factors related to the presence of the parent tree could be affecting the quality of the soil if each species uses nutrients in a particular way. In this scenario, the soil in the vicinity of a parent tree would have a lower quantity of particular nutrients necessary for the seedlings favouring dense or negative distance-dependent effects on seedling recruitment. If this were to happen, the diversity of neotropical forests could also be promoted by distance-dependent abiotic processes. To explore these processes, we tested the following hypotheses and predictions:

1. Since the establishment of seedlings depends on the amount of nutrients available (Clark et al., 1998), adding fertilizer should increase seedling’s survival and growth.
2. Competition with tree roots affects the performance of understory plants (Wright 2002, Wright et al., 2015), seedlings that are isolated from the parent’s roots should perform better.
3. Negative density-dependent processes can be explained by abiotic effects (i.e., depletion of particular nutrients used by the parent, Wills et al., 1997), seedlings planted in soil collected away from the parent trees should have better growth and survival.
4. Some plant species accumulate pathogens rapidly and maintain low densities due to the negative effect of species-specific pathogens (Klironomos, 2002, Bagchi et al 2014). Thus, seedlings planted in soil collected far from the parent trees should have a better development since potential species-specific pathogens would not be as abundant. We expected a similar effect for seedlings planted in parent soil treated with Agrodyne®, a bactericide-fungicide.
5. Competition for light with canopy plants affects the performance of plants in the understory (Wright, 2002, Dormann et al., 2020). Seedlings planted in the shade house - located in a forest clearing with greater light availability-, should have higher growth than those growing in the forest understory.

## Methods

This study was carried out at the El Zafire Biological Station in the Colombian Amazon at 4°0’21”S and 69°53’55” W (Figure 1). We used seedlings from the trees *Caraipa* sp. (Clusiaceae) and *Tachigali* sp. (Fabaceae), for which we located 10 seedling banks (i.e., 10 parent trees) and *Dicranostyles* sp. (Convolvulaceae) for which eight seedling banks were found.

**Figure 1.**
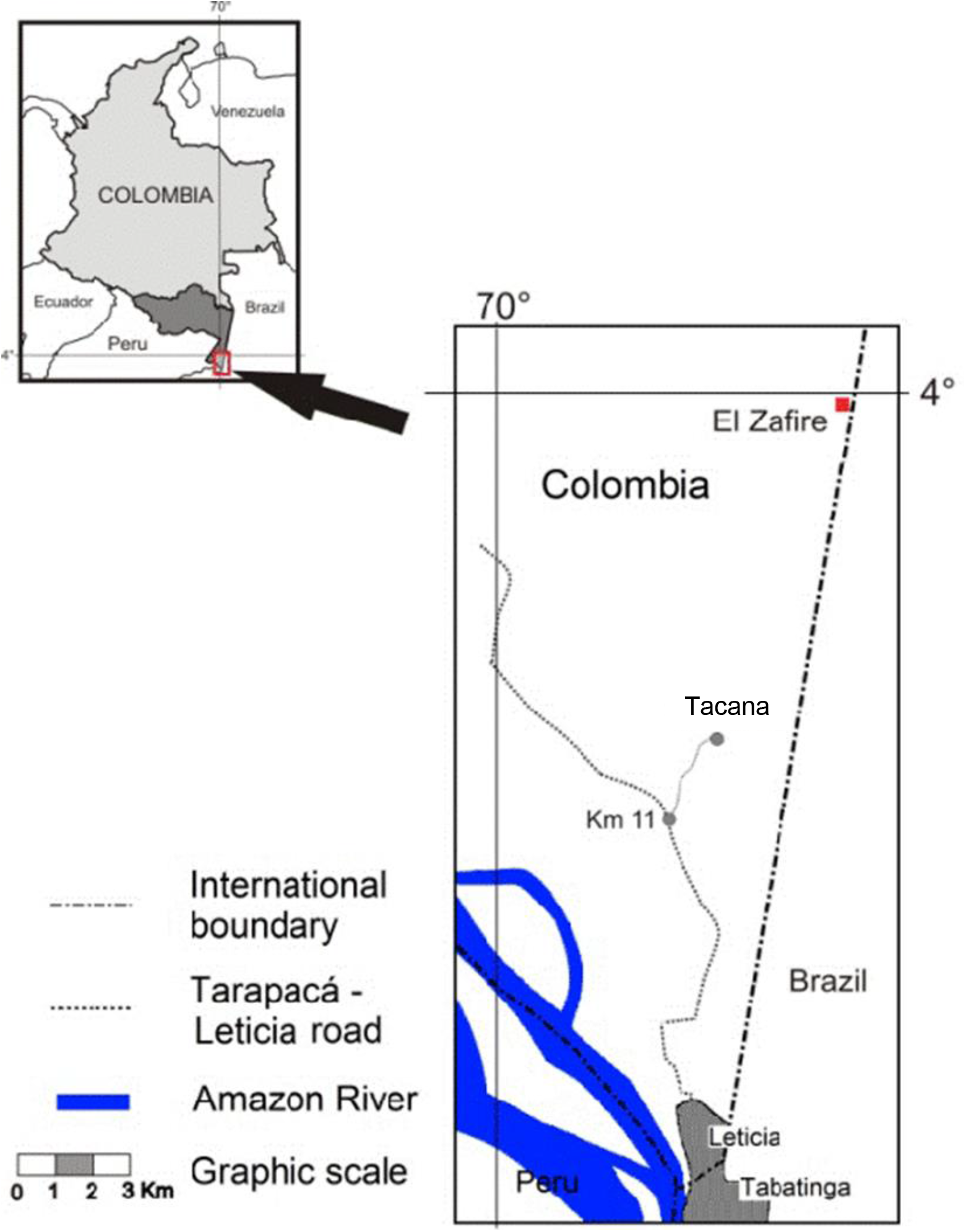
Study area, El Zafire Biological Station. Figure taken from Navarro et al. 2011.

To apply the different treatments we planted groups of 10 seedlings per treatment in the forest and in the shade house.

### Forest experiment

Under each parent tree we randomly allocated 10 seedlings to each of the following treatments:

1. Added fertilizer (hypothesis 1)
2. Excluded from the parental roots (hypothesis 2)
3. Planted on soil collected far from the parent tree (hypotheses 3 and 4).
4. Control: unearthed and replanted seedlings to account for the manipulation of all seedlings.

### Shade house experiment

We planted 30 seedlings per parent tree. Seedlings were randomly allocated to the following treatments:

1. Planted on soil collected around the parent tree (hypotheses 3 and 4).
2. Planted on soil collected around the parent tree treated with Agrodyne ^®^ (hypotheses 3 and 4).
3. Planted on soil collected far from the parent tree. As a control to evaluate the effect of the previous two treatments and to compare the performance of plants exposed to more light (Hypothesis 5).

### Measurements

To calculate final growth, we used the relative growth rate (RGR) calculated in mm per day from the difference between initial and final heights of each seedling divided by the growth period (Baraloto et al., 2005).

We determined five potential causes of seedling death:1) Herbivory; 2) Falling debris; 3) Added fertilizer; 4) Stem breakage; and 5) Death for no apparent reason.

We measured light intensity using a 401025 Ex Tech Light meter on 10 clear sky days in an open clearing, the understory and the shade house.

We assessed the effects of the different treatments on the growth of our seedlings using ANOVA models (control vs treatment) and analysed survivorship for both experiments using the “survival” (Therneau 2020) and “survminer” (Kassambara et al., 2021) R packages. We analysed pairwise comparisons using Log-Rank Test (p value adjustment method for multiple comparisons: BH). Analyses were performed using R (R Development Core Team, Version 1.1.1335).

## Results

### Forest experiment

The relative growth rate (RGR) of our study species was not significantly affected by the addition of fertilizer. Isolating seedlings from the parent’s roots increased the RGR of *Caraipa* seedlings, slowed the RGR for *Tachigali* and had no effect on *Dicranostyles*. Control seedlings of *Caraipa* had a higher RGR than seedlings growing on soil collected away from the parent tree while *Dicranostyles* and *Tachigali* seedlings were not affected by this treatment (Figures 2a, 2b and 2c).

**Figure 2.**
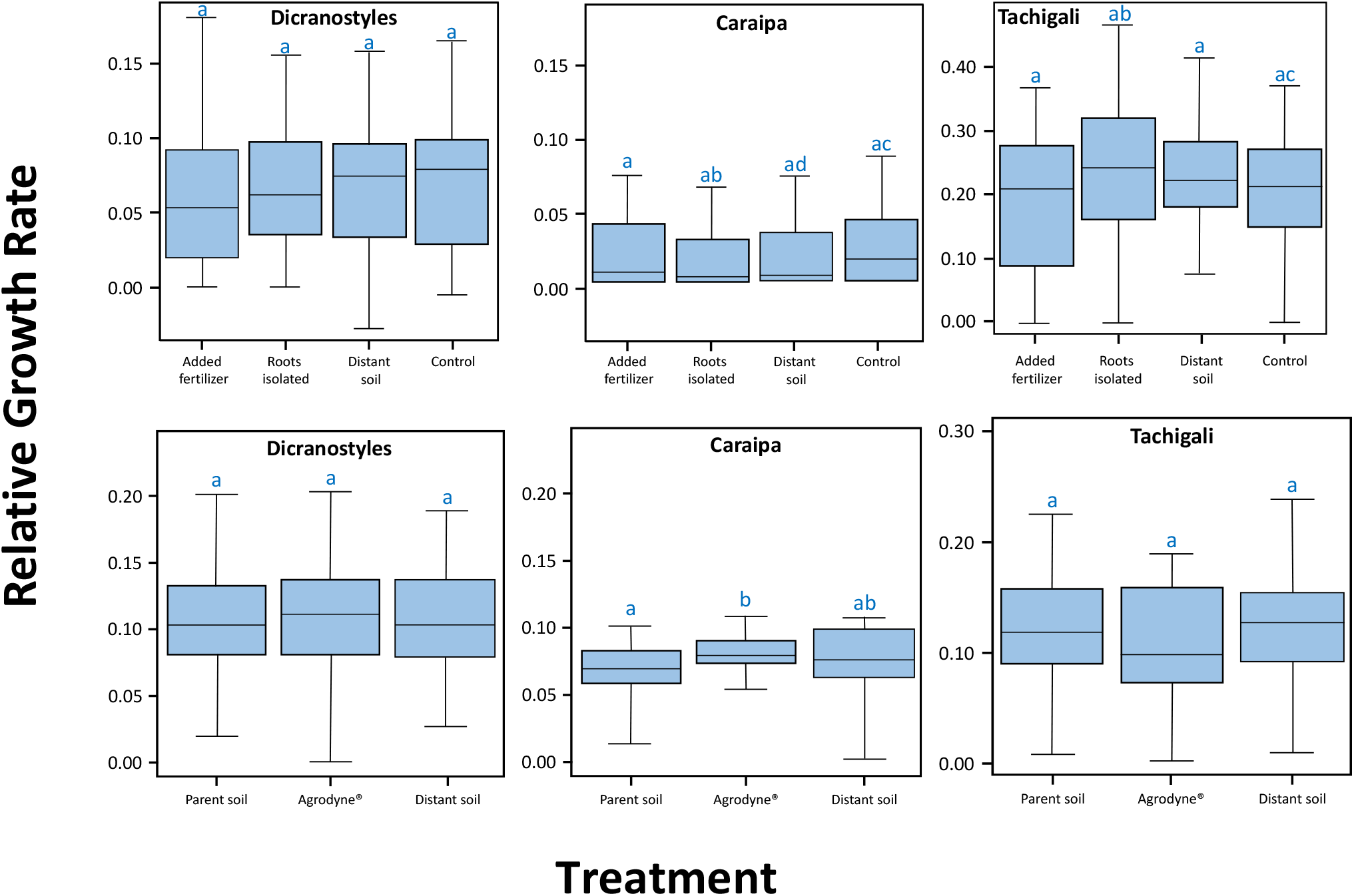
Growth of seedlings for all species and treatments. The top row corresponds to seedlings growing in the forest, bottom row corresponds to seedlings growing in the shade house. Different letters indicate significant difference between treatments. The tick line represents the median, the outer limits of the box the first and third quartiles. Whiskers extend to cover any data point <1.5 times the interquartile range.

Survival of all seedlings planted in the forest ranged between 97.5% and 24%. Seedlings of *Caraipa* growing isolated from neighbouring roots had the highest survival (97.5%). The lowest survival was recorded for *Tachigali* seedlings growing with addition of fertilizer (24%) (Figures 3a, 3b and 3c).

**Figure 3.**
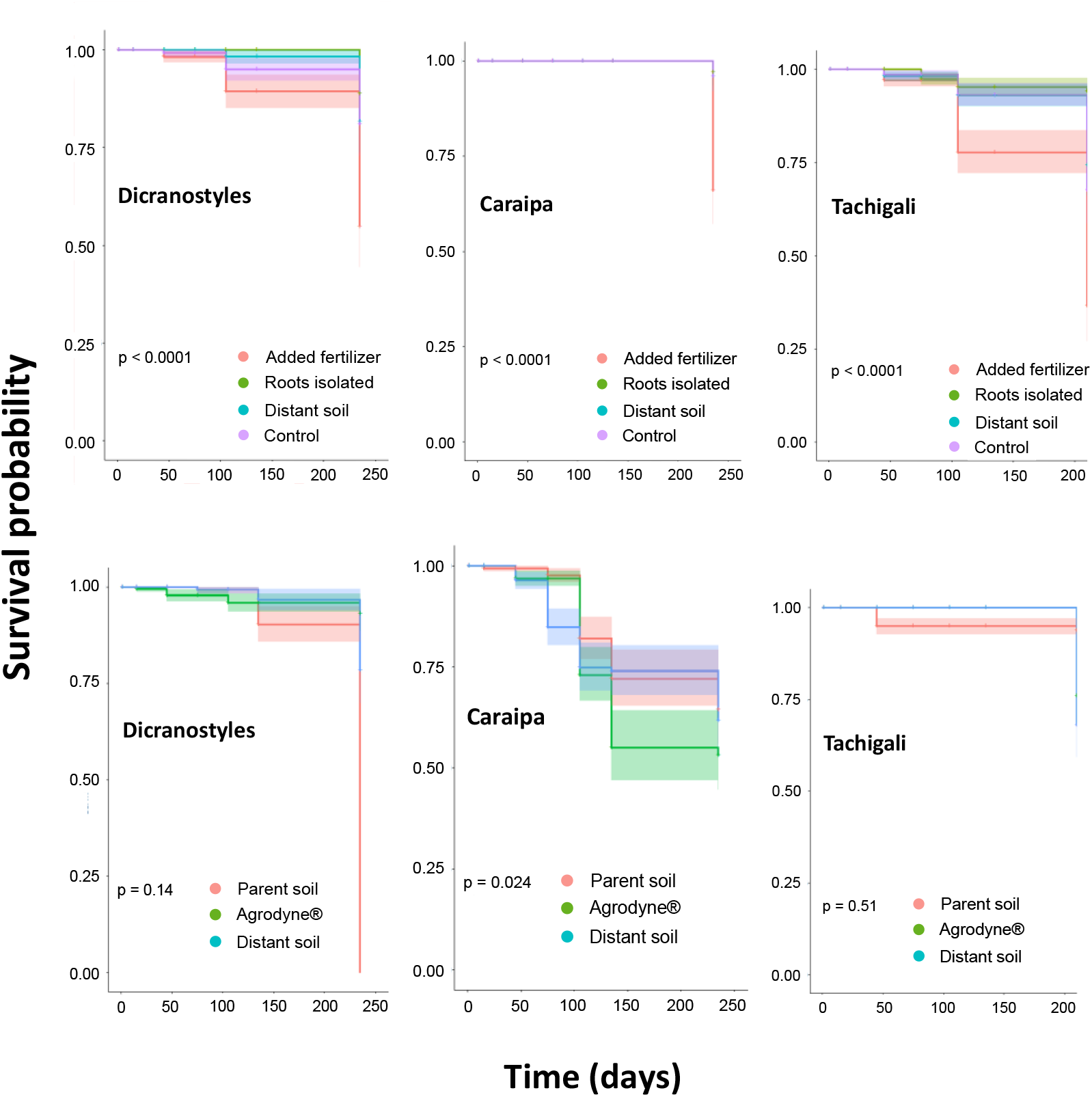
Survival of planted seedlings. The top row corresponds to seedlings planted in the forest; bottom row corresponds to seedlings planted in the shade house.

### Shade house experiment

We found significant differences between the RGR of *Caraipa* and *Tachigali* seedlings planted on soil treated with Agrodyne^®^ compared to seedlings planted on soil collected near the parent tree; the effect was different for each species. *Caraipa* seedlings planted on soil treated with Agrodyne^®^ had a higher RGR than seedlings planted on soil collected near the parent tree, the opposite pattern was recorded for *Tachigali* (Figure 2e and 2f).

Survival of seedlings planted in the shade house ranged between 98% and 28.7%. The highest survival was recorded for seedlings of *Tachigali* planted on soil collected near the parent tree (98%), the lowest survival was for *Caraipa* seedlings planted on soil collected far from the parent tree (35%). (Figures 3d, 3e and 3f).

#### Light availability

The amount of light that reaches the understory and the shade house was 2.6% and 51.5%, respectively compared to the open clearing (100%). For all species the seedlings planted in the shade house showed a higher RGR than seedlings planted in the forest (Figure 2).

#### Overall mortality

The causes of mortality for seedlings planted in the shade house were difficult to determine, dead seedlings showed no obvious signs of herbivory, fungi, or any other damage. The causes of mortality for seedlings planted in the forest were quantified, for all species the addition of fertilizer caused the highest mortality (Table 1).

**Table 1.** Mortality causes of seedlings growing in the forest

| **Number of death seedlings per specie** | **Addition of fertilizer** | **Fallen branches or other debris** | **No apparent reason** | **Herbivory** | **Broken stem** |
| --- | --- | --- | --- | --- | --- |
| *Caraipa* sp – 48 | 62.5 % | 0% | 33.3% | 4.2% | 0 |
| *Tachigali* sp – 182 | 19.8% | 27.6% | 3.2% | 46.2% | 3.2% |
| *Dicranostyles* sp - 67 | 10.5% | 10.5% | 32.8% | 0% | 0 |

## Conclusions

Our results provide evidence of the lack of a simple and common response to the studied factors across species. These findings indicate that the response of tropical seedling communities to local abiotic and biotic factors are highly dynamic and species specific.

We found two common responses across species: we recorded higher growth rates for seedlings planted in the shade house where more light was available; this emphasizes the importance of light as a limiting factor for seedling growth in tropical forests (Montgomery & Chazdon 2002, Umaña et al. 2018). The other common response was the high mortality recorded across species when fertilizer was added, and its lack of effect on plant growth. Other studies have found similar patterns including higher mortality of individuals with added nutrients due to increase competition (Norden et al. 2007), or no effect of nutrients on plant performance (Mangan et al. 2010).

We found contrasting results for the three species of seedlings replanted in the forest. For example, seedlings from *Caraipa* sp showed differences in their RGR between the control and the seedlings isolated from other roots, and seedlings growing in soil collected far from the parent tree. However, contrary to what we expected, those with the highest RGR were the control seedlings. This may be because both groups of seedlings: isolated from other roots, and seedlings planted in soil collected far from the parent tree were planted in bags, reducing their available area. Seedlings with a smaller available area grow less and are more likely to die than those with a larger available area (Mithen et al. 1984). For seedlings of *Tachigali* sp we also found significant differences between the seedlings isolated from neighbouring roots and the control. In this case, the results were as we expected, seedlings isolated from other roots grew more.

We found distinct responses for all seedlings planted in the shade house. Seedlings of *Tachigali* sp showed significant differences between the RGR of seedlings treated with Agrodyne^®^ and those that were planted in soil collected far from the parent tree. Contrary to what we expected and to the hypothesis of species-specific pathogen associations, the seedlings treated with Agrodyne^®^ grew less than the others. It is likely that the untreated soil contains favourable microorganisms that promote germination and establishment, as has been found in other studies (Sarmiento et al. 2017). The vital role of soil microbial communities in maintaining rainforest diversity has been highlighted by many authors (e.g., Mangan et al. 2010; Comita et al. 2014, Bever et al. 2015 and Wood et al. 2020, among others); therefore, it is not surprising that reducing or eliminating the soil microorganisms could have a negative effect on the growth of certain seedlings.

In our study, each species responded differently to the studied factors, preventing us to draw general conclusions that could help explain the high plant diversity of tropical moist forests. It should be noted that the species we studied were selected due to their abundance. These three species were the only ones with eight or more large seedling banks; therefore, it is possible that these species are good competitors, and that for this reason they are abundant and able to tolerate different conditions without showing major changes in their growth and survival.

Detailed methods and results are provided as supplementary information.

## Acknowledgements

We would like to thank Maria Cristina Peñuela and Adriana Aguilar for their support to carry out our research at El Zafire Biological Station. Edilberto, Ángel, Jairo and Magnolia provided field assistance built the shade house. Eufrasia helped locating the seedlings banks.

## Financial support

This research received no specific grant from any funding agency, commercial or not-for-profit sectors.

## Competing interests

The authors declare none.

## Data Availability Statement

The data that support the findings of this study are available upon request

## Notes

### Competing Interest Statement

The authors have declared no competing interest.

