## supplementary information for "Exploring the role of abiotic factors on the seedling recruitment of three plant species in the Colombian Amazon"

**Supplementary information – Detailed methods and results**

### Methods

#### *Study area*

This study was carried out at the El Zafire Biological Station located in the Calderón River Forest Reserve, Colombian Amazon at 4°0'21"S and 69°53'55" W (Figure 1). The forest surrounding the research station includes three forest types: white sands, temporarily flooded, and *terra firme*. Our study was carried out in *terra firme* forest.

#### *Species used*

To select our species, we scanned the study site looking for extensive seedling banks of recently germinated seedlings. We found three plant species with seedling banks large enough to setup all experiments; the trees: *Caraipa* sp. (Clusiaceae) and *Tachigali* sp. (Fabaceae), for which we located 10 seedling banks (i.e., 10 parent trees and *Dicranostyles* sp. (Convolvulaceae) for which eight seedling banks (eight parent trees) were found.

To apply the different treatments we planted groups of 10 seedlings per treatment in the forest and in the shade house; with 10 or eight replications per treatment, depending on the number of parent trees found (i.e., 10 for *Caraipa* sp and *Tachigali* sp. and eight for *Dicranostyles* sp.).

#### *Forest experiment*

In our field experiment we used four treatments to test the different hypotheses. Under each parent tree we randomly allocated seedlings to each of the following treatments:

1. Fertilizer addition: to evaluate hypothesis 1 (nutrient-limited recruitment).

2. Excluded from the parental roots: to evaluate hypothesis 2 (competition for nutrients limits the growth and survival of seedlings).

3. Planted on soil collected far from the parent tree: to evaluate the effect of distance-dependent processes of abiotic and biotic nature (since the microorganisms in the area can colonize the experimental stations).

4. Controls: to evaluate recruitment without any treatment. We unearthed and replanted these seedlings to account for the manipulation of the other seedlings.

In the forest we planted 40 seedlings per parent tree, out of these we added NPK fertilizer to 10 seedlings to test whether competition for nutrients limits their growth and/or survival. We added fertilizer at the beginning of the experiment, three weeks later, and then once per month until the end of the experiment. We isolated other 10 seedlings from the surrounding roots and planted them in bags to avoid the rapid inclusion of roots from the parent and other nearby trees and seedlings. Additionally, we planted 10 seedlings on soil collected away from the parent trees. These seedlings were also planted in plastic bags to prevent this soil from mixing with the local soil. Finally, we replanted 10 seedlings as a control, under the parent tree as the other seedlings, and replanted to account for the manipulation of all other seedlings.

***Shade house experiment***

In our shade house experiment we used three treatments to test the hypotheses. We randomly allocated seedlings to each of the following treatments:

1 Seedlings planted on soil collected around the parent tree: to evaluate recruitment with potential distance-dependent abiotic and biotic effects.

2. Seedlings planted on soil collected around the parent tree treated with Agrodyne ® (bactericide-fungicide): to evaluate potential distance-dependent abiotic effects.

3. Seedlings planted on soil collected far from the parent tree. As a control to evaluate the effect of the previous two treatments and to compare the performance of plants exposed to more light.

We planted 30 seedlings per parent tree. We planted the seedlings in plastic bags in groups of 10. To evaluate recruitment with potential distance-dependent biotic and abiotic effects we planted seedlings on soil collected around the parent tree (10 per tree). For our second treatment, we soaked the soil collected around the parent tree with Agrodyne®, using a solution of 5 ml per litre (as per product instructions). We immersed the stems and roots of the seedlings planted in this treated soil in an Agrodyne® solution for 5 minutes (5 ml per liter). Finally, we planted seedlings in soil collected far from the parent tree; this soil was collected 50 metres after the area occupied by the seedling banks had ended. Since at 50 metres from the parent tree distance-dependent processes are not usually evident (Hubbel et al. 2001, Murphy *et al*., 2017). Thus, we could assume that this soil was already far enough to avoid any effect of the parent tree and/or the seedling bank.

**Seedling measurements**

***Growth***

After all seedlings were replanted we recorded their initial height. Two weeks later we measured them again and then every month during four months for *Dicranostyle*s sp. and *Caraipa* sp. Seedlings of *Tachigali* sp. were measured every month for three months. Final measurements were taken seven months after the first measurement for *Dicranostyles* sp and *Caraipa* sp and after six months for *Tachigali* sp. To calculate the final growth of the seedlings we used the relative growth rate (RGR) (Baraloto et al., 2005). We calculated RGR in mm per day from the difference between initial and final heights of each seedling divided by the growth period ((ln (height final) − ln (height initial))/ (growth period in days)). The total height of each seedling was measured from the soil to the most distant part of the main stem.

***Mortality***

We determined five potential causes of seedling death:1) Death by herbivory, when the seedling was clearly eaten. 2) Death due to falling leaf litter, logs and/or branches. 3) Death by added fertilizer, since the seedlings that suffered the greatest mortality in the forest were those treated with fertilizer; at first these seedlings showed black leaves and later died. Although this type of mortality is not common, in another study with seedlings and addition of nutrients the same pattern emerged (Norden et al., 2007). 4) Death due to stem breakage, and 5) Death for no apparent reason, when the seedling died with no obvious signs of any type of damage.

***Light availability***

To assess whether light availability had a significant impact in the growth and survival of seedlings we measured light intensity using a 401025 Ex Tech Light meter. We measured light intensity on 10 clear sky days in an open clearing, in the understory and in the shade house.

***Data analysis***

We assessed the effects of the different treatments on the growth of our seedlings using ANOVA models (control vs treatment) and analysed survivorship for both experiments using the “survival” (Therneau 2020) and “survminer” (Kassambra et al. 2019) R packages. We analysed pairwise comparisons using Log-Rank Test (p value adjustment method for multiple comparisons: BH). Analyses were performed using R (R Development Core Team, Version 1.1.1335). All data were normal, except the growth data for the seedlings of *Tachigali* sp growing in forest, these data were transformed using the root of the tangent to obtain normality.

### Results

#### ***Forest experiment***

*Relative growth rate across treatments*

Contrary to what we expected, the relative growth rate (RGR) of our study species was not significantly altered by the addition of fertilizer (*Dicranostyles*, F = 0.28, p = 0.60; *Caraipa*, F = 0.45, p = 0.50; *Tachigali*, F = 0.04, p = 0.84). However, isolating seedlings from the parent's roots increased the RGR of *Caraipa* seedlings (F = 0.75, p = 0.03);we found the opposite effect for *Tachigali* (F = 5.45, p = 0.02), and no effect on *Dicranostyles* (F = 0.28, p = 0.60). Unexpectedly, the control seedlings of *Caraipa* (i.e., planted around the parent tree, on the same soil), showed a higher RGR than the seedlings growing on soil collected away from the parent tree (F = 3.95, p = 0.04), while *Dicranostyles* and *Tachigali* seedlings were not affected by this treatment (F = 0.02, p = 0.96 and F = 2.48, p = 0.11, respectively) (Figures 2a, 2b and 2c).

*Survival*

Survival of seedlings planted in the forest ranged between 97.5% and 24%. Seedlings of *Caraipa* showed higher survival, with seedlings growing isolated from neighbouring roots showing the highest survival (97.5%), followed closely by seedlings growing on soil collected far from the parent tree and control seedlings (96.5% and 96.1%, respectively). We recorded the lowest survival for this species on seedlings growing with addition of fertilizer (66.55%, p < 0.0001 for all comparisons). For *Dicranostyles* seedlings growing isolated from neighbouring roots had the highest survival (89.5%), followed by seedlings growing on soil collected far from the parent tree (79 %) and control seedlings (72.7%). We also recorded the lowest survival for *Dycranostyles* for seedlings growing with addition of fertilizer (44.5%, p < 0.0001 for all comparisons). We recorded the lowest survival for *Tachigali*, for which the highest survival was found for seedlings growing on soil collected far from the parent tree

(63.5%), followed closely by seedlings growing isolated from neighbouring roots (63%) and control seedlings (58%). Once again, we recorded the lowest survival for this species for seedlings growing with addition of fertilizer (24%, $p < 0.0001$ for all comparisons) which also was the lowest survival registered for all seedlings planted in the forest (Figures 3a, 3b and 3c).

#### *Shade house experiment*

*Relative growth rate across treatments*

For seedlings growing in the shade house, we found significant differences between the RGR of *Caraipa* and *Tachigali* seedlings planted on soil treated with Agrodyne® compared to seedlings planted on soil collected near the parent tree; again the effect was different for each species. For *Caraipa*, seedlings planted on soil treated with Agrodyne® had a higher RGR than seedlings planted on soil collected near the parent tree ($F=4.64$, $p=0.03$), we recorded the opposite pattern for *Tachigali* ($F=6.21$, $p=0.01$) (Figure 2e and 2f). We did not find any significant differences in the RGR between seedlings planted on soil collected far from the parent tree and seedlings planted on soil collected near the parent tree (*Dicranostyle*s: $F=0.25$, $p=0.65$; *Caraipa*: $F=1.14$, $p=0.28$ and *Tachigali*: $F= 0.08$, $p= 0.77$) (Figures 2d, 2e and 2f).

*Survival*

Survival of seedlings planted in the shade house ranged between 98% and 28.7%. We recorded the highest survival for seedlings of *Tachigali* planted on soil collected near the parent tree (98%), followed by seedlings planted on soil treated with Agrodyne® (95%). The lowest survival for this species was recorded for seedlings planted on soil collected far from the parent tree (86%). For *Dicranostyles*, we recorded the highest survival for seedlings planted on soil treated with Agrodyne® (84%), followed by seedlings growing on soil collected near the parent tree (83%). The lowest survival recorded for this species was for seedlings growing on soil collected far from the parent tree (80%,). In *Caraipa*, we found the highest survival for seedlings planted on soil collected near the parent tree (42%), followed by seedlings planted on soil collected far from the parent tree (35%). We recorded the lowest survival for this species for seedlings planted on soil treated with Agrodyne® (28.7%) (Figures 3d, 3e and 3f).

*Light availability*

We recorded the highest light availability in the open clearing (1,156 ± 30 lux, n = 10), followed by the shade house (595 ± 24 lux, n = 10). The understory had the lowest light availability (30 ± 0 lux, n = 10). The amount of light that reaches the understory and the shade house was 2.6% and 51.5%, respectively compared to the open clearing (100%).

As expected, for all species the seedlings planted in the shade house showed a higher RGR than seedlings planted in the forest (*Caraipa* sp, $F = 124.28$, $p \leq 0.001$; *Tachigali* sp, $F = 49.47$, $p \leq 0.001$; *Dicranostyles* sp., $F = 171.28$, $p \leq 0.001$; Figure 2).

*Overall mortality*

The causes of mortality for seedlings planted in the shade house were difficult to determine, dead seedlings showed no obvious signs of herbivory, fungi, or any other damage. It is possible that differences in humidity and temperature between the shade house and the forest (i.e., hotter and drier conditions) may have increased plant mortality in the shade house. The causes of mortality for seedlings planted in the forest were quantified, as mentioned before for all species the addition of fertilizer caused the highest mortality.
